# PEDF/PEDF-R Signaling Regulates Retinal Phospholipid Homeostasis and Photoreceptor Survival

**DOI:** 10.64898/2026.08.04.742541

**Authors:** Alexandra Bernardo-Colón, Susan E Crawford, Martin Paul Agbaga, Zhen Wang, Kevin Schey, S. Patricia Becerra

**Affiliations:** Section of Protein Structure and Function, LRCMB, NEI-NIH; Department of Cancer Biology, Endeavor Health Research Institute, Affiliate of University of Chicago Pritzker School of Medicine, Evanston, IL, USA; Department of Cell Biology, Department of Ophthalmology, and Dean A. McGee Eye Institute, Oklahoma City, OK, USA; Department of Biochemistry, Vanderbilt University School of Medicine

## Abstract

Pigment epithelium-derived factor (PEDF) promotes photoreceptor survival through its receptor PEDF-R, a phospholipase involved in retinal lipid metabolism. To define the in vivo function of the PEDF/PEDF-R axis, we generated mice lacking *Serpinf1* (PEDF) and *Pnpla2* (PEDF-R). Combined loss of *Serpinf1* and *Pnpla2* resulted in severe retinal degeneration characterized by outer nuclear layer (ONL) thinning, outer segment (OS) shortening, reduced rhodopsin and cone opsin expression, increased TUNEL-positive nuclei, and enhanced retinal autofluorescence associated with altered lipid distribution. Lipid-associated markers, including TIP47, PLIN5, and BODIPY, exhibited abnormal distribution patterns in mutant retinas, indicating disrupted lipid storage and trafficking. Loss of PEDF/PEDF-R signaling also impaired photoreceptor-rod bipolar cell connectivity, as demonstrated by reduced PKCα/synaptophysin colocalization, and resulted in diminished electroretinographic responses. Lipid Imaging mass spectrometry revealed decreases in some lipid abundances in photoreceptor outer segment and inner segment/outer nucleus layer, while lipids containing arachidonic acid and docosahexaenoic acid-containing lipids showed increased abundance. Together these findings identify the PEDF/PEDF-R signaling axis as a key regulator of retinal phospholipid homeostasis that couples lipid metabolism to photoreceptor survival and visual function.

## Introduction

Photoreceptors maintain one of the most extensive and dynamic membrane systems in the body, continuously renewing phospholipid-rich outer segment discs that are essential for phototransduction and vision (1). Maintaining phospholipid homeostasis—the regulated balance of membrane phospholipid composition, distribution, and turnover—is therefore essential for photoreceptor integrity and visual function. Consequently, disruption of phospholipid homeostasis can compromise photoreceptor survival and contribute to retinal degeneration, a leading cause of blindness worldwide.

Pigment epithelium-derived factor (PEDF), encoded by *SERPINF1*, is a neurotrophic factor secreted by the retinal pigment epithelium—the monolayer of cells behind the photoreceptors—that promotes photoreceptor survival through PEDF-R, encoded by *PNPLA2* (2–4). PEDF-R also known as ATGL, *PNPLA2* or desnutrin is a patatin-like domain phospholipase that has been studied extensively for its triglyceride lipase activity in adipose tissue but is thought to function predominantly as a phospholipase in the phospholipid-rich retina (2). Upon PEDF binding, PEDF-R-mediated PLA₂ activity hydrolyzes membrane phospholipids to release fatty acids and lysophospholipids implicated in cell survival and regeneration (2, 3, 5, 6). Photoreceptor outer segments are enriched in DHA-containing phospholipids. and undergo continual membrane renewal, creating a sustained demand for phospholipid remodeling to preserve membrane integrity (7). PEDF-R phospholipase activity is therefore well positioned to contribute to the phospholipid remodeling required for photoreceptor membrane renewal.

Ablation of *Pnpla2* in mice causes photoreceptor malformation, impaired phagocytosis, and reduced visual function (8–10). Consistent with a role for PEDF-R in retinal phospholipid metabolism, loss of *Pnpla2* is associated with accumulation of lysophospholipid species, including LysoPC-DHA and LysoPE-DHA, indicating disruption of phospholipid remodeling pathways in the retina (8). In contrast, *Serpinf1* null mice exhibit no overt retinal or photoreceptor abnormalities under baseline conditions but display increased susceptibility to stress-associated retinal degeneration (11). Together, these findings implicate the PEDF/PEDF-R signaling axis in photoreceptor maintenance and retinal phospholipid homeostasis, yet its physiological role in coordinating these processes in vivo remains undefined. To address this question, we generated mice lacking both PEDF (*Serpinf1*) and PEDF-R (*Pnpla2*) to define the role of the PEDF/PEDF-R signaling axis in retinal phospholipid homeostasis, photoreceptor integrity, and visual function.

## Results

### Loss of *Serpinf1* and *Pnpla2* expression in the retina and retinal pigment epithelium

To investigate the role of the PEDF/PEDF-R axis in retinal lipid remodeling and photoreceptor integrity *in vivo*, we analyzed mice deficient in *Serpinf1* and *Pnpla2*. Because constitutive *Pnpla2* (*PEDF-R*) knockout mice develop systemic lipid abnormalities and die by 3–4 months of age, likely due to cardiac lipid accumulation (Fig. S1), all experiments were performed in animals younger than 3 months. To confirm depletion of *Serpinf1* and *Pnpla2* in the retina, transcript levels were measured by qRT-PCR using whole-retina RNA. *Serpinf1* transcripts were reduced by approximately 50% in *Serpinf1^+/-^*;*Pnpla2^-/-^* mice and were undetectable in *Serpinf1^-/-^;Pnpla2^-/-^* retinas (Fig. S2A). Similarly, *Pnpla2* transcripts were undetectable in both *Serpinf1^+/-^* ;*Pnpla2^-/-^ and Serpinf1^-/-^;Pnpla2^-/-^* retinas (Fig. S2B), confirming efficient depletion of both genes.

Because the RPE is the principal source of retinal PEDF, we next examined PEDF protein levels in RPE extracts. PEDF protein was readily detected in wild-type (*Serpinf1*^+/+^;*Pnpla2*^+/+^) mice at 3 months of age (Fig. S2C). In contrast, PEDF protein levels were reduced by approximately 40% in *Serpinf1^+/-^* ;*Pnpla2^-/-^* and markedly reduced (83% reduction) in *Serpinf1^-/-^;Pnpla2^-/-^ mice (*Fig. S2C).

Together, these results confirm depletion of *Serpinf1*/PEDF and *Pnpla2*/PEDF-R in the retina and establish the genetic models used for subsequent analysis.

### PEDF/PEDF-R deficiency causes retinal structural abnormalities and photoreceptor degeneration

To determine the consequences of PEDF/PEDF-R deficiency on retinal structure, we examined retinal cross-sections from mutant and control mice. Compared with wild-type littermates, both *Serpinf1^-/-^;Pnpla2^-/-^* and *Serpinf1^+/-^;Pnpla2^-/-^* mice exhibited marked thinning of the outer nuclear layer (ONL) (Fig. 1A). Although all major retinal layers were present across genotypes, overall retinal thickness decreased with increasing loss of both PEDF and PEDF-R. Quantitative analysis of ONL thickness at defined dorsal and ventral retinal positions surrounding the optic nerve head was visualized as a radial (spider) plot (Fig. 1B). This analysis demonstrated a consistent reduction in ONL thickness across both retinal regions in *Serpinf1^-/-^;Pnpla2^-/-^* and *Serpinf1^+/-^; Pnpla2^-/-^* mice compared with *wild-type* controls.

**Figure 1:**
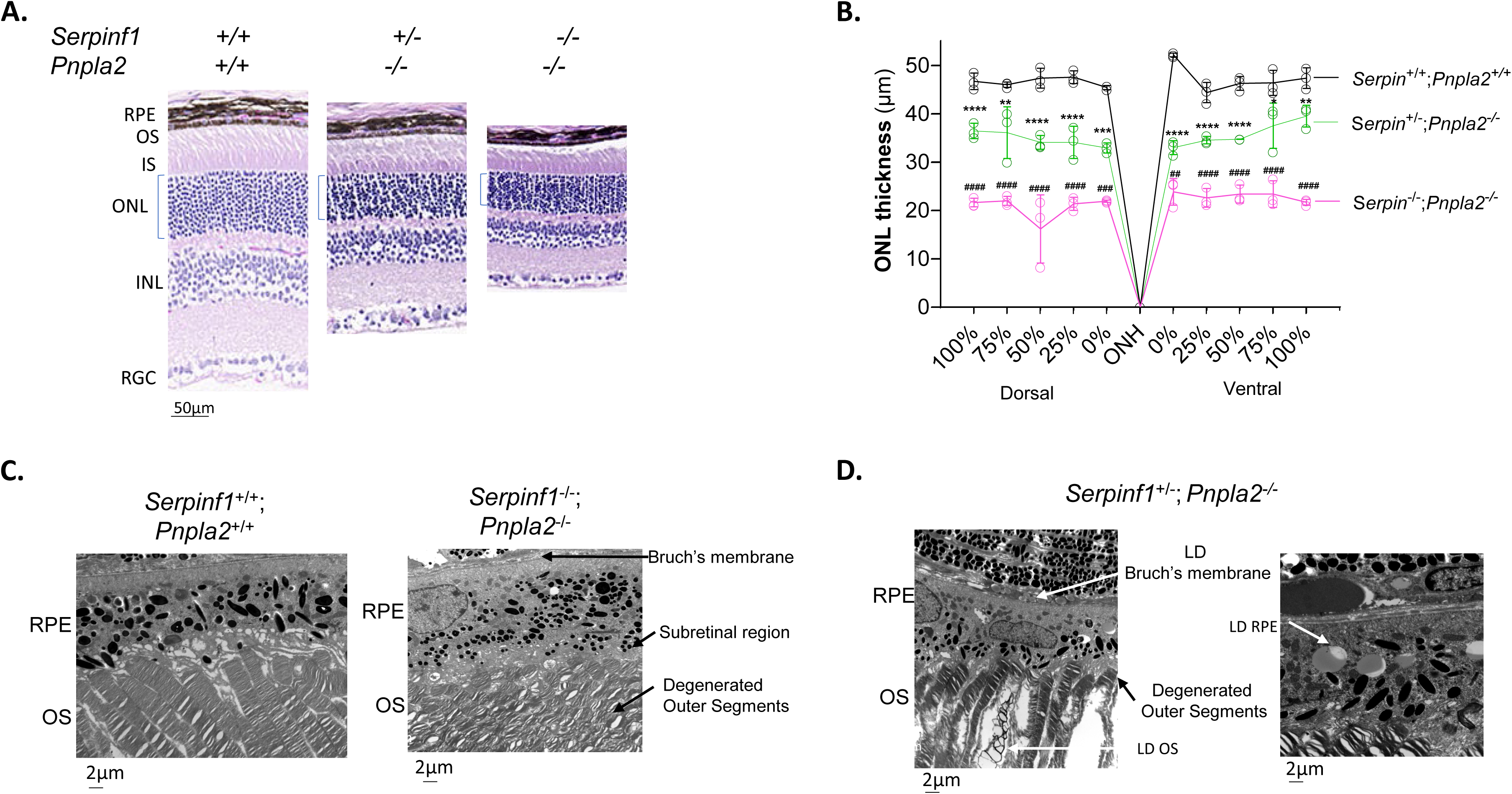
Histological and ultrastructural evaluation of PEDF/PEDF-R depletion in the retina and retinal pigment epithelium (RPE) of the *Serpinf1/Pnpla2* mouse line. (A) Microphotographs of retina sections of *Serpinf1^+/+^;Pnpla2^+/+^*, *Serpinf1^+/-^*;*Pnpla2^-/-^* and *Serpinf1^-/-^*;*Pnpla2^-/-^* mice at 3 months of age, stained with hematoxylin and eosin (B) Quantification of outer nuclear layer (ONL) thickness across dorsal and ventral regions relative to the optic nerve head presented as a spider plot. Five retinas per group were evaluated, and each data point corresponds to the average ± SD per location. Statistical analysis was performed using one-way ANOVA with Dunette’s multiple comparisons test. ∗*P* < 0.05, ∗∗*P* < 0.01, ∗∗∗*P* < 0.001, ∗∗∗∗*P* < 0.0001 and for *Serpinf1^+/-^*/*Pnpla2^-/-^* ##*P* < 0.01, ###*P* < 0.001, ####*P* < 0.0001 relative to *Serpinf1^+/+^*/*Pnpla2^+/+^* mice. Scale bar=50µm. (C) Representative transmission electron microscopy (TEM) images of retinal and RPE ultrastructure from *Serpinf1^+/+^* x *Pnpla2^+/+^*, *Serpinf1^+/-^* x *Pnpla2^-/-^*and *Serpinf1^-/-^* x *Pnpla2^-/-^* mice at 3 months of age. Images were selected from four eyes of each genotype (*n* = 4). Scale bar: is 2μm.

To further assess retinal ultrastructure, retinas from *Serpinf1*^+/-^;*Pnpla2*^-/-^, Serpinf*1*^-/-^;*Pnpla2*^-/-^ and wild-type littermates were examined by transmission electron microscopy (TEM). In wild-type mice, RPE cells showed organized structure with intact apical processes and basal infoldings (Fig. 1C). The subretinal space between the photoreceptor outer segments and the RPE contained electron-dense material consistent with lipid-containing material. Photoreceptor outer segments were well organized, with regularly spaced discs, and their apical tips were tightly ensheathed by the RPE apical processes. In *Serpinf1*^+/-^;*Pnpla2*^-/-^ mice (Fig. 1D), the Bruch’s membrane appeared largely intact but was thickened and showed early degenerative changes. Numerous lipid droplets accumulated within the RPE cytoplasm, Bruch’s membrane, and choriocapillaris (black arrows), suggesting altered lipid handling within the outer retina. Although overall RPE organization was relatively preserved, distal photoreceptor outer segments showed membrane disruption and disc disorganization, accompanied by lipid accumulation between adjacent outer segments. In contrast, *Serpinf1*^-/-^;*Pnpla2*^-/-^ mice (Fig. 1C) exhibited severe ultrastructure abnormalities. Photoreceptor outer segments were highly disorganized, with malformed and degenerated membranes and discs. RPE cells appeared thickened, with disorganized and amorphous apical processes that failed to properly ensheathe outer segment. This was associated with a markedly reduced subretinal space, characterized by densely packed outer segments and reduced extracellular material (subretinal region). The Bruch’s membrane was further thickened and showed pronounced degenerative changes.

### PEDF/PEDF-R deficiency reduces photoreceptor visual pigment expression

Rhodopsin and cone opsins are visual pigment proteins localized to photoreceptor outer segment discs, where they are embedded within phospholipid-rich membranes and mediate phototransduction (12, 13). To assess the effects of PEDF/PEDF-R deficiency on photoreceptor integrity, we examine visual pigment expression by immunofluorescence.

Compared with wild-type littermate controls, both *Serpinf1^-/-^;Pnpla2^-/-^* and *Serpinf1^+/-^;Pnpla2^-/-^* mice exhibited markedly reduced rhodopsin (55% and 64% reductions, respectively) and opsin (61% and 66% reductions, respectively) immunoreactivity (Fig. 2A). Quantitative analysis showed that rhodopsin and opsin levels were reduced to approximately 35-40% of wild-type levels in both mutant genotypes (Figs. 2B and 2C).

**Figure 2:**
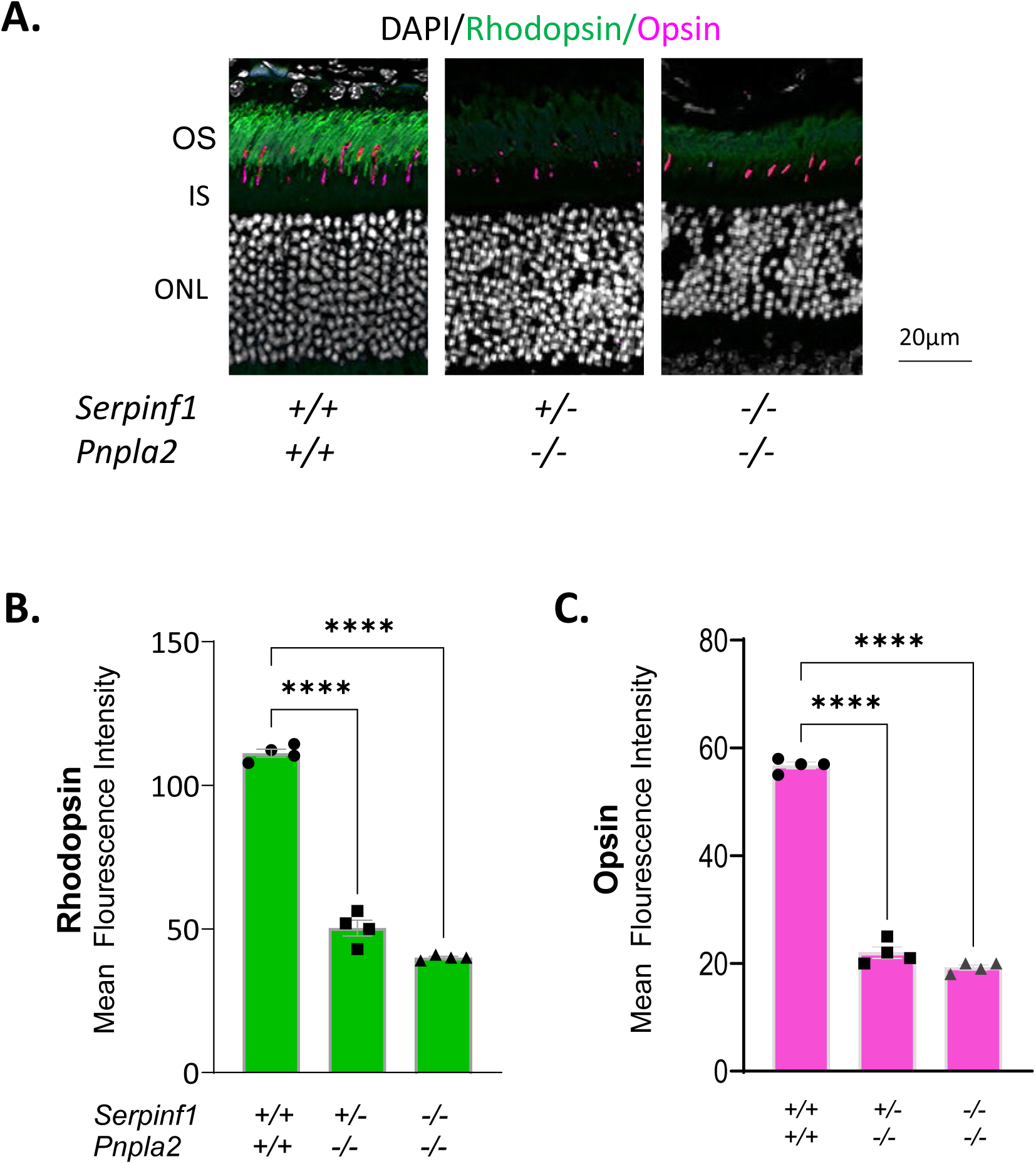
Reduced photoreceptor marker immunoreactivity in PEDF/PEDF-R–deficient retinas. (A) Representative immunofluorescence images of retinal sections stained for rhodopsin (rods, green), cone opsin (magenta), and nuclei (DAPI, blue) from *Serpinf1^+/+^*;*Pnpla2^+/+^* (WT), *Serpinf1^+/−^*;*Pnpla2^−/−^*, and *Serpinf1^−/−^*;*Pnpla2^−/−^* mice at 3 months of age. (B, C) Quantification of rhodopsin and cone opsin fluorescence intensity. Four retinas per genotype (n = 4) were analyzed. For each retina, 3–5 fields were imaged and averaged to obtain a single value. Data are presented as mean ± SD. Statistical analysis was performed using one-way ANOVA with Dunnett’s multiple comparisons test versus WT. ****P < 0.0001. Scale bar, 20 μm.

These results demonstrate that combined deficiency of PEDF and PEDF-R reduces visual pigment abundance in photoreceptors.

### Photoreceptor cell death in PEDF/PEDF-R-deficient mice

To assess the impact of PEDF/PEDF-R deficiency on photoreceptor survival, retinal cell death was evaluated using PSVue-488, a fluorescent probe that binds phosphatidylserine (PS) exposed on the outer leaflet of the plasma membrane during early stages of cell death (14, 15). PSVue-488 was administered topically as eye drops to *Serpinf1^-/-^;Pnpla2^-/-^*, *Serpinf1^+/-^;Pnpla2^-/-^*, and wild-type mice. Fundus fluorescence imaging was performed 24 h after administration to detect surface-exposed PS.

As shown in Fig. 3A, retinas of *Serpinf1^+/-^;Pnpla2^-/-^* mice exhibited increased fluorescence intensity relative to contralateral eyes treated with vehicle alone (HBSS). The fluorescence signal was further increased in *Serpinf1^-/-^;Pnpla2^-/-^* mice, whereas wild-type controls displayed minimal labeling. Quantitative analysis confirmed significantly higher fluorescence intensity in both mutant genotypes relative to controls (Fig. 3B), indicating enhanced PS exposure in PEDF/PEDF-R-deficient retinas.

**Figure 3:**
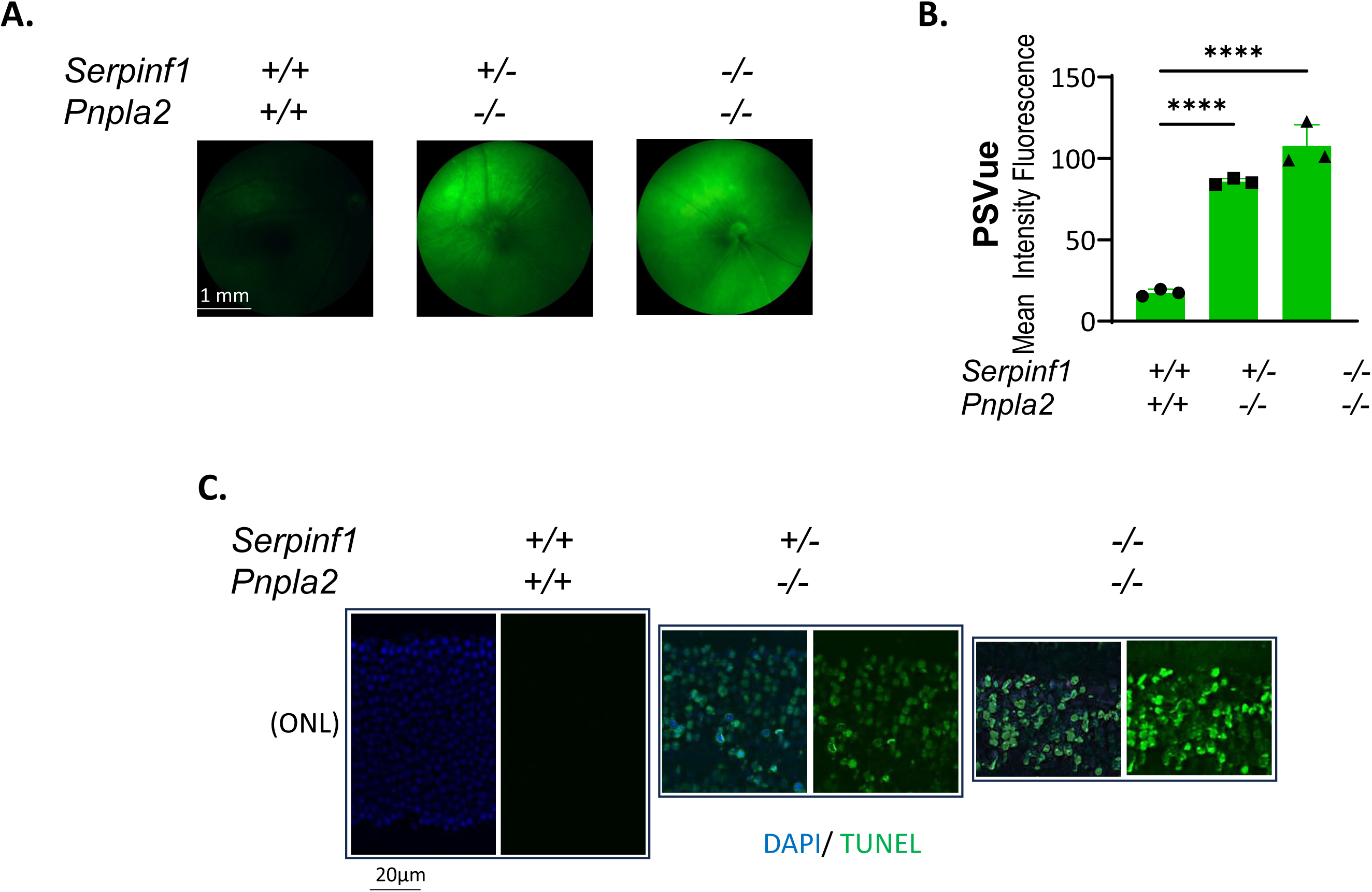
Increased photoreceptor cell death in PEDF/PEDF-R–deficient mice. (A) Representative fundus fluorescence images showing PSVue-488 labeling of phosphatidylserine exposure in retinas from *Serpinf1*^+/+^;*Pnpla2*^+/+^ (WT), *Serpinf1*^+/−^;*Pnpla2*^−/−^, and *Serpinf1*^−/−^;*Pnpla2*^−/−^ mice. Control eyes treated with HBSS alone were used for background subtraction. (B) Quantification of PSVue-488 fluorescence intensity. (C) Representative retinal sections stained for TUNEL (green) and DAPI (blue). Three retinas per genotype (n = 3) were analyzed. Data are presented as mean ± SD. Statistical analysis was performed using one-way ANOVA with Dunnett’s multiple comparisons test versus WT. \*\**P* < 0.001, \*\*\**P* < 0.0001. Scale bar, 20 μm.

To determine the retinal localization of the fluorescence signal eyes from PSVue® 488-treated *Serpinf1^+/-^*;*Pnpla2^-/-^* and *Serpinf1^-/-^;Pnpla2^-/-^* mice were collected 24 h after administration and processed for fluorescence microscopy. As shown in Fig. S3, PSVue® 488 labeling (green signal indicated by white arrows) was localized to the outer nuclear layer (ONL), as well as in the outer plexiform layer (OPL)indicated by white arrows consistent with phosphatidylserine externalization in photoreceptor cells undergoing degeneration. These findings indicate that the fluorescence signal detected by fundus imaging originates primarily from the photoreceptor layer.

To further evaluate consequence of PEDF/PEDF-R-deficiency on photoreceptor cell survival or death, retinal sections were analyzed by TUNEL staining. TUNEL-positive nuclei were rarely observed in the ONL of wild-type retinas but were readily detected in both *Serpinf1^+/-^;Pnpla2^-/-^* and *Serpinf1^-/-^;Pnpla2^-/-^* mice (Fig. 3C). In *Serpinf1^-/-^;Pnpla2^-/-^* retinas, the ONL nuclei exhibited pronounced pyknotic morphology consistent with advanced stages of cell death.

Together, these findings demonstrate that combined loss of PEDF and PEDF-R is associated with increased photoreceptor cell death in the retina, which supports the loss of retinal ONL and significant reduction in total retinal thickness.

### Altered lipid distribution and lipid droplet organization in PEDF/PEDF-R-deficient retinas

To assess alterations in lipid handling associated with PEDF/PEDF-R deficiency, we examined the localization of lipid-droplet-associated proteins and neutral lipids in the retina by immunofluorescence. We first analyzed TIP47 (tail-interacting protein of 47 kDa) that interacts with lipid droplets (16). In wild-type retinas, TIP47 was predominantly localized to photoreceptor outer segments (OS) tips and the RPE (Fig. 4A indicated by white arrows). In both *Serpinf1^+/-^;Pnpla2^-/-^* and *Serpinf1^-/-^;Pnpla2^-/-^*mice, TIP47 immunoreactivity remained evident at OS tips and within the RPE and choroid (Fig. 4A). Notably, in *Serpinf1^+/-^;Pnpla2^-/-^* retinas, TIP47 signal was also detected within the ONL suggesting altered intracellular localization in photoreceptors. A similar pattern of TIP47 staining, localized primarily to the RPE/choroid and photoreceptor outer segments (OS), was observed in *Pnpla2^-/-^* mice. In contrast, little to no TIP47 immunoreactivity was detected in *Serpinf*1^-/*-*^ (Fig. S4A).

**Figure 4:**
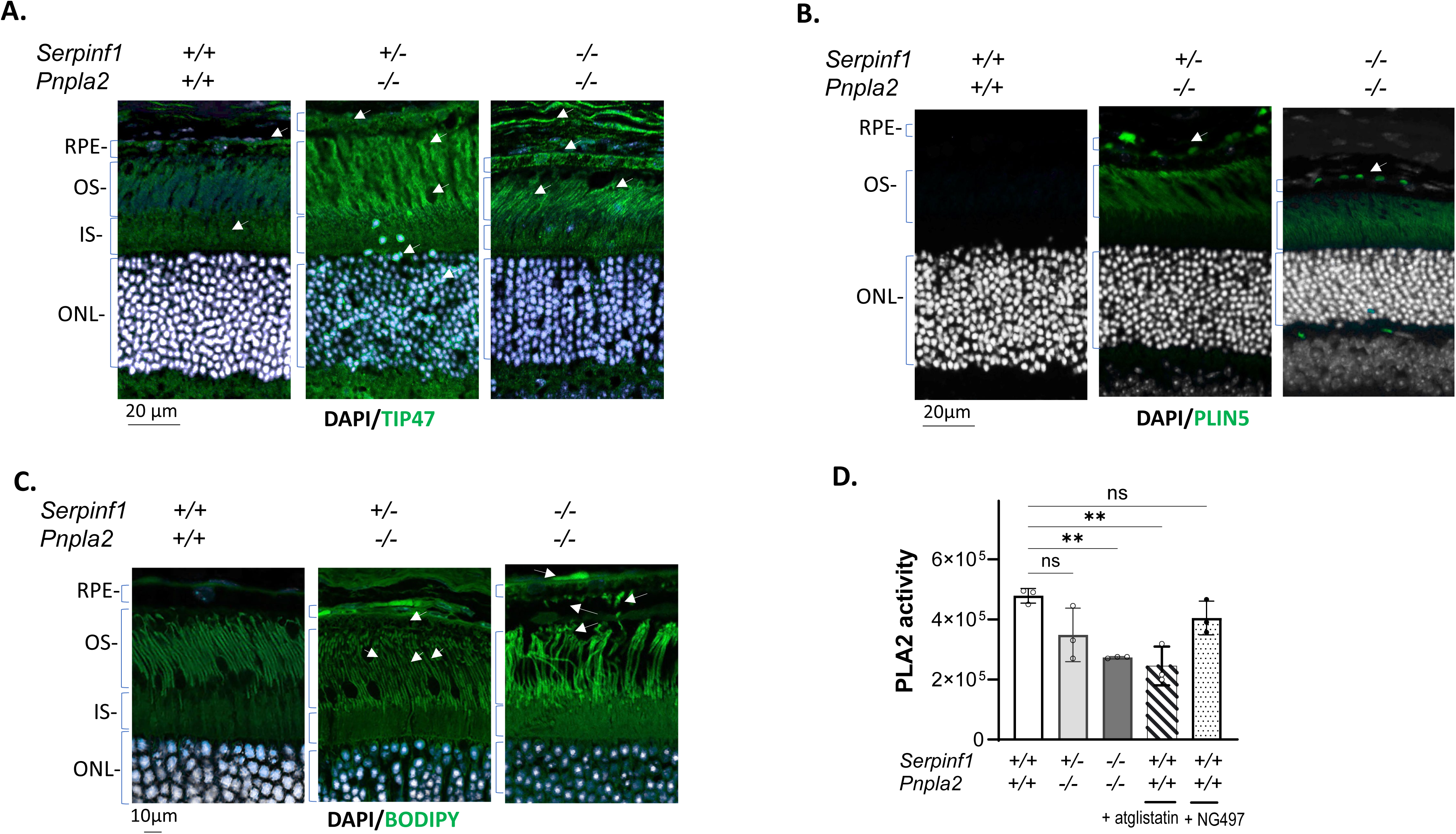
Lipid accumulation and altered lipid-associated protein localization in PEDF/PEDF-R–deficient retinas. (A-C) Representative fluorescence micrographs of retinal sections from *Serpinf1^+/+^*;*Pnpla2^+/+^*, *Serpinf1^+/-^*;*Pnpla2^-/-^* and *Serpinf1^-/-^*;*Pnpla2^-/-^*, mice at 3 months of age stained for TIP47, PLIN5 or BODIPY 493/503. (A) TIP47 immunofluorescence in photoreceptors outer segments (OS), outer nuclear layer (ONL) and RPE. (B) PLIN5 immunofluorescence in the RPE and photoreceptor OS tips. White arrows indicate PLIN5-positive puncta. (C) BODIPY 493/503 staining shows lipids within photoreceptors and the RPE; white arrows indicate lipid deposits. For all immunofluorescence analyses, three retinas per genotype were analyzed (n=3). Scale bars, 10 μm (C) and 20μm (A-B). (D) Retinal phospholipase A2 (PLA2) activity measured retinal extracts from *Serpinf1^+/+^*;*Pnpla2^+/+^*, *Serpinf1^+/-^*;*Pnpla2^-/-^* and *Serpinf1^-/-^*;*Pnpla2^-/-^* mice at 3 months of age. Wild-type retinal extracts were incubated with 3.5 μM atglistatin (hatched bar), or NG-497 (patterned bar). Each data point represents a single retina, and the bars correspond to the mean ± SD (n = 3 per genotype). Statistical analysis was performed using one-way ANOVA with Dunnett’s multiple comparisons test. \**P* < 0.05.

Next, we examined perilipin 5 (PLIN5), a lipid droplet–associated protein involved in lipid storage and utilization through interactions between lipid droplets and mitochondria (17). Compared with wild-type controls, both mutant genotypes exhibited increased PLIN5-associated lipid droplets in the RPE (Fig. 4B indicated by white arrows). In *Serpinf1^+/-^; Pnpla2^-/-^* and *Serpinf1^-/-^;Pnpla2^-/-^* mice, PLIN5 immunoreactivity appeared as discrete puncta throughout the basal side of the RPE and choroid and OS tips. Consistent with the TIP47, *Pnpla2^-/-^ retinas exhibited* strong PLIN5 staining in the RPE, indicative of lipid droplet accumulation. In contrast, *Serpinf1^-/-^ r*etinas showed little to no detectable PLIN5 staining, suggesting that PEDF deficiency alone does not promote lipid droplet accumulation in the RPE (Fig. S4B). This observation suggests that PEDF deficiency alone alters PLIN5 distribution differently from combined PEDF/PEDF-R deficiency.

To directly visualize neutral lipid accumulation, retinal sections were stained with BODIPY 493/503 (18). BODIPY labeling revealed increased neutral lipid droplets within the RPE and focal lipid deposits at photoreceptor OS tips in both S*erpinf1^+/-^;Pnpla2^-/-^* and *Serpinf1^-/-^;Pnpla2^/-^*mice relative to wild-type controls (Fig. 4C).

Together, these findings reveal genotype-dependent alterations in retinal lipid organization, including altered localization of lipid-associated proteins and accumulation of neutral lipid deposits in PEDF/PEDF-R-deficient mice.

### PEDF-R contribution to retinal PLA2 activity

To assess the contribution of PEDF-R to total PLA₂ activity, wild-type retinal extracts were incubated with the inhibitors atglistatin and NG-497. Atglistatin reduced PLA₂ activity, whereas NG-497 had no detectable effect under the assay conditions used (Fig. 4D). Residual PLA₂ activity was observed in both PEDF/PEDF-R–deficient retinas and atglistatin-treated wild-type extracts, suggesting that additional retinal PLA₂ enzymes likely contribute to the measured activity. Together, these findings indicate that total retinal PLA₂ activity reflects the combined contribution of multiple phospholipases and suggest that PEDF-R contributes to retinal PLA₂ activity.

### Imaging mass spectrometry and lipid profiling of PEDF/PEDF-R-deficient retinas

To further characterize the lipid alterations observed by immunofluorescence, we performed lipid profiling and imaging mass spectrometry (IMS) analysis of retinas from wild-type, *Serpinf1*^+/−^;*Pnpla2*^−/−^, and *Serpinf1*^−/−^;*Pnpla2*^−/−^ mice. These analyses were performed to characterize genotype-dependent changes in retinal lipid composition and spatial lipid distribution associated with PEDF/PEDF-R deficiency.

IMS analysis of retina sections from each genotype (WT, Serpinf1+/−;Pnpla2−/−, and Serpinf1−/−;Pnpla2−/−) generated heatmaps containing approximately 150 signals in positive ion mode and 200 signals in negative ion mode. The majority of these signals were identified as lipids and did not exhibit changes in either intensity or spatial distribution. However, intensity changes were detected for numerous lipid species, with the affected signals predominantly localized to the photoreceptor outer segment (OS) or the inner segment/outer nuclear layer (IS/ONL). Overlays of selected lipids across different retina layers in both positive mode and negative mode are shown in Figure 5. Consistent with photoreceptor degeneration and thinning of the ONL observed in other analyses, several lipids showed reduced abundance in the photoreceptor and ONL regions (Fig. 5A, 5A′, and 5B′), including PI 16:0_20:5 detected in negative mode and PC 14:0_16:0 detected in positive mode.

**Figure 5:**
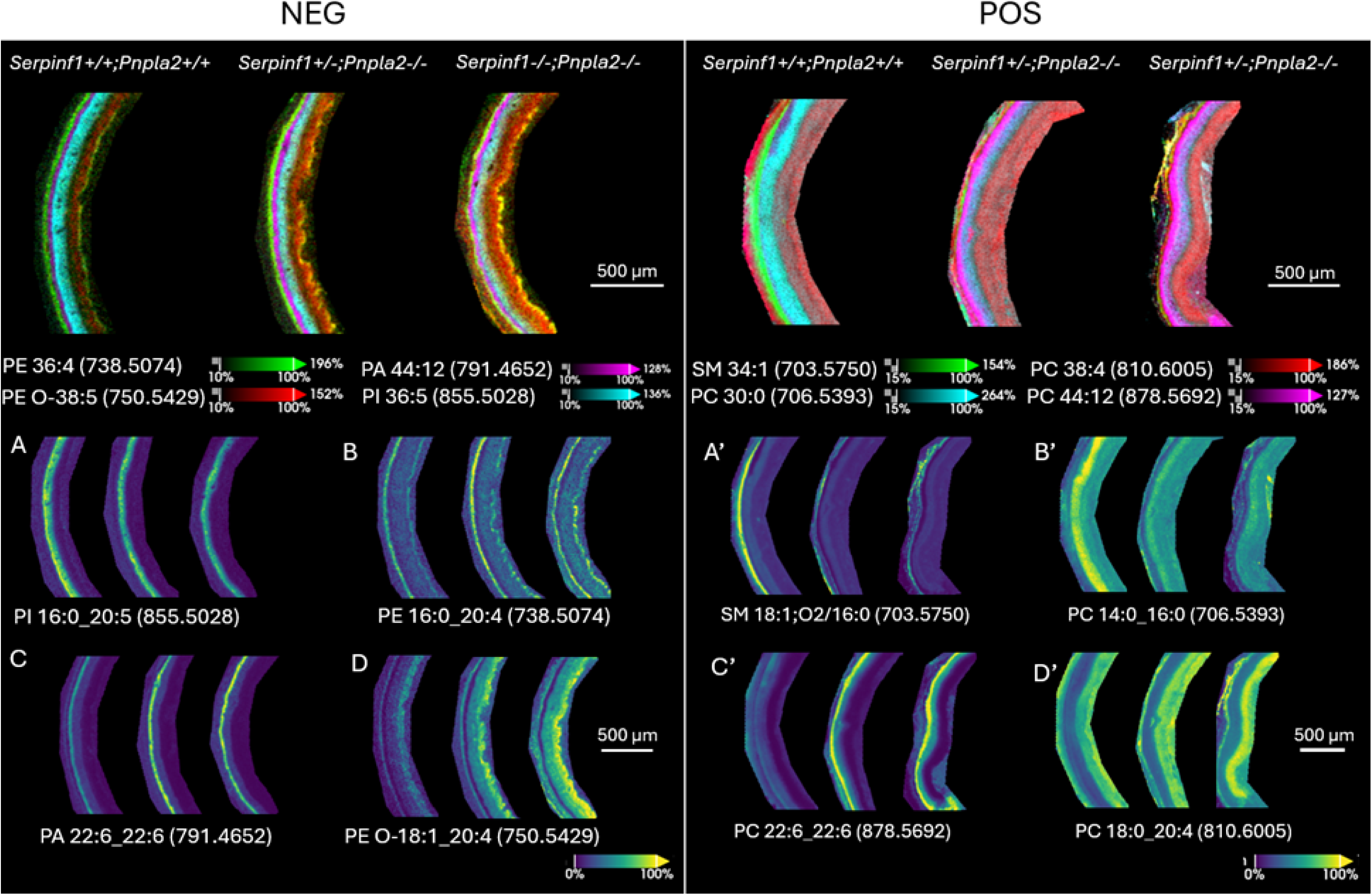
MSI analysis of lipids in WT and PEDF/PEDF-R-deficient mice. Overlays of four lipid signals (top panels) in both negative and positive mode reveal lipids distributed in specific retinal layers and abundance changes in *Serpinf1^+/-^;Pnpla2^-/-^, and Serpinf1^-/-^;Pnpla2^-/-^* mice retina. Images for individual lipid signals are shown below (A-D for negative data and A’-D’ for positive data). All lipids shown were identified by LC-MS/MS. Scale bars: 500 μm.

In contrast, several arachidonic acid-containing lipids, including PE 16:0_20:4 (Fig. 5B), PE O-18:0_20:4 (Fig. 5D), and PC 18:0_20:4, were significantly increased in the photoreceptor region and the inner retina. These lipids were primarily localized to the RPE/choroid or the inner retinal layers. In addition, several DHA-containing lipids, including PA 22:_22:6 and PC 22:6_22:6, localized to the photoreceptor outer segments also showed increased abundance, as illustrated in Figs. 5C and 5C′, respectively.

These findings indicate that loss of PEDF/PEDF-R alters retinal phospholipid composition and spatial lipid organization, particularly within photoreceptor-associated regions.

### Altered retinal function and rod bipolar cell dendritic remodeling in PEDF/PEDF-R-deficient mice

Given the structural abnormalities observed in photoreceptors, retinal function was assessed by electroretinography (ERG). Representative ERG traces for S*erpinf1^+/-^;Pnpla2^-/-^* and *Serpinf1^-/-^;Pnpla2^-/-^* and wild-type mice are shown in Fig. 6A. Quantitative analysis revealed that maximal a-wave amplitudes (a_max) were significantly reduced in both mutant genotypes relative to wild-type mice controls (Fig. 6B). Similarly, maximal b-wave amplitudes (b_max) were markedly decreased, consistent with impaired transmission of photoreceptor-driven signals to downstream bipolar cells.

**Figure 6:**
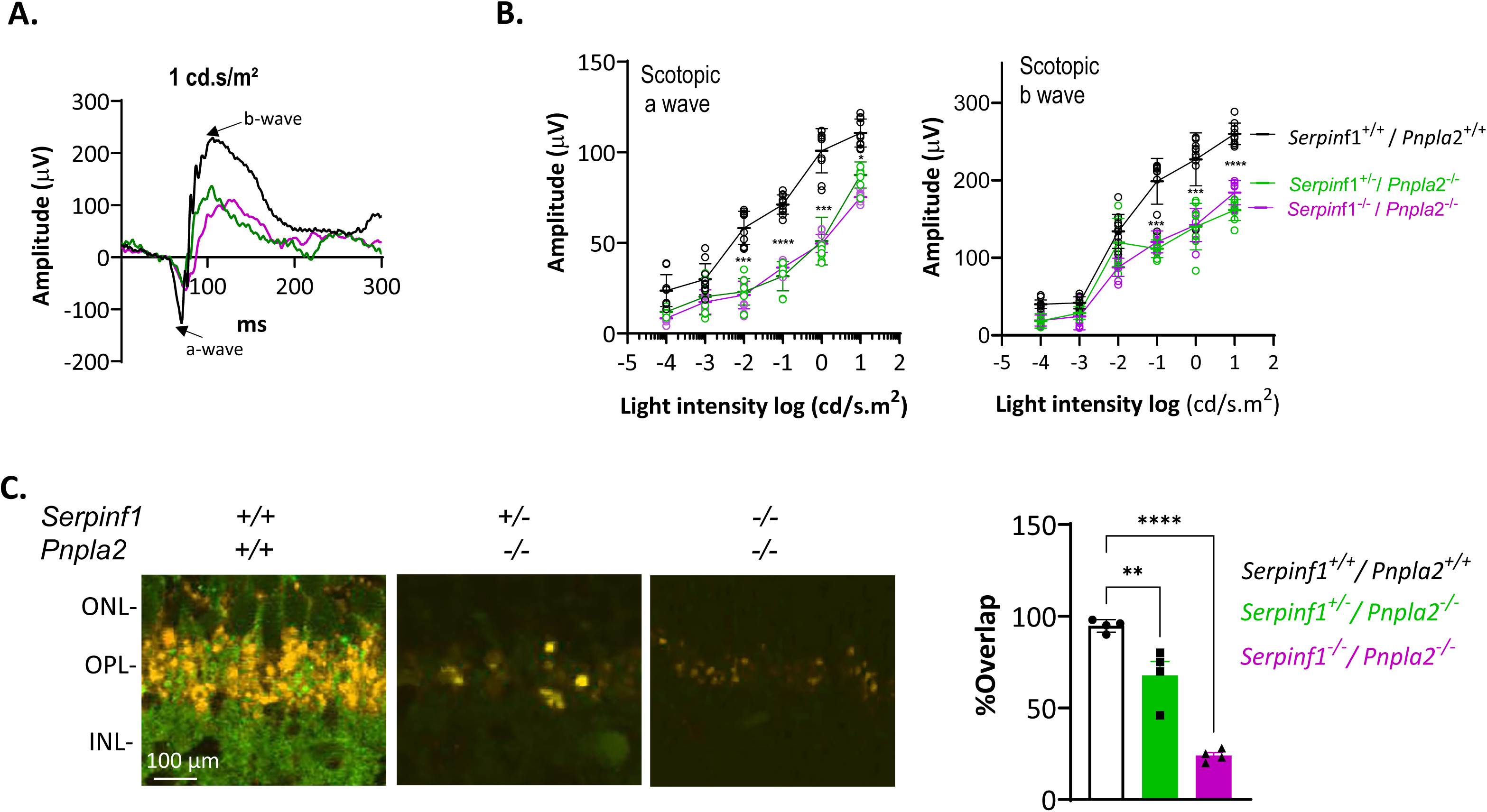
Impaired retinal function and synaptic connectivity in PEDF/PEDF-R–deficient mice. (A) Representative scotopic ERG traces from *Serpinf1*^+/+^;*Pnpla2*^+/+^ (WT), *Serpinf1*^+/−^;*Pnpla2*^−/−^, and *Serpinf1*^−/−^;*Pnpla2*^−/−^ mice at 3 months of age (1 cd·s/m² flash). (B) Quantification of a-wave and b-wave amplitudes as a function of light intensity. Each data point represents one mouse (WT, n = 10; *Serpinf1*^+/−^;*Pnpla2*^−/−^, n = 8; *Serpinf1*^−/−^;*Pnpla2*^−/−^, n = 10). Values from both eyes were averaged for each mouse. Data are presented as mean ± SD. (C) Representative immunofluorescence images of retinal sections stained for PKCα (rod bipolar cells, green) and synaptophysin (presynaptic terminals, red), with quantification of fluorescence signal overlap. Scale bar, 100 μm. Four retinas per genotype (n = 4) were analyzed for immunofluorescence. Data are presented as mean ± SD Statistical analysis was performed using one-way ANOVA with Dunnett’s multiple comparisons test versus WT. \**P* < 0.05, \*\**P < 0.01, ***P < 0.001, ****P<0.0001*.

To further assess synaptic retraction in the associated with the reduced b-wave responses, rod bipolar cell dendrites were examined by PKCα immunolabeling together with the presynaptic marker synaptophysin. In wild-type retinas, rod bipolar dendrites extended throughout the outer plexiform layer and showed substantial overlap with synaptophysin-positive terminals (Figs. 6C). In contrast, *Serpinf1^+/-^;Pnpla2^-/-^* and *Serpinf1^-/-^;Pnpla2^-/-^* retinas exhibited reduced overlap between PKCα and synaptophysin labeling, consistent with altered rod bipolar cell dendritic organization and reduced interaction with photoreceptor terminals (Fig. 6C). Quantitative colocalization analysis confirmed a significant overlap reduction in both mutant genotypes relative to controls, consistent with the observed ERG deficits.

Together, these results indicate that loss of PEDF and PEDF-R impairs photoreceptor-driven responses and is associated with altered synaptic organization between photoreceptors and rod bipolar cells at the first visual synapse.

## Discussion

Our study demonstrates that double-knockout PEDF/PEDF-R mice exhibit reduced outer nuclear layer (ONL) thickness, disorganized photoreceptor outer segments (OS), decreased rhodopsin and opsin levels, synaptic retraction in the outer plexiform layer (OPL), and compromised visual function. These structural and functional deficits are accompanied by increased photoreceptor cell death and dysregulated lipid accumulation. Notably, retinal degeneration was substantially more severe in homozygous double knockout mice than previously reported for PEDF-R deficiency alone (8), indicating that ligand-receptor disruption produces defects beyond loss of receptor function alone. Together, these findings demonstrate that PEDF/PEDF-R signaling is essential for maintaining photoreceptor integrity, retinal phospholipid homeostasis and visual function. PEDF is increasingly recognized for its neurotrophic functions in the retina, mediated specifically through interaction of its neurotrophic domain with PEDF-R (2, 19, 20). In our study, no obvious vascular abnormalities were observed among the genotypes (Fig. S5), suggesting that the retinal degeneration primarily reflects disruption of neurotrophic and metabolic signaling rather than vascular pathology. Previous studies showed that PEDF deficiency sensitizes photoreceptors to stress and promotes neovascularization without causing overt degeneration (11, 21), supporting a model in which PEDF sustains PEDF-R-dependent neuroprotective signaling in the retina.

PEDF-R (encoded by *PNPLA2*) has been extensively studied for its triglyceride lipase activity in peripheral tissues, but it exhibits phospholipase activity in the retina and represents an important component of retinal phospholipid metabolism. Loss of PEDF-R alone leads to retinal degeneration (8, 10), while simultaneous PEDF/PEDF-R deficiency produces even more severe abnormalities, emphasizing the cooperative role of ligand-receptor signaling. Phospholipids, particularly those enriched in DHA and arachidonic acid (AA), are critical for photoreceptor membrane integrity, energy metabolism, and signaling (22, 23). Imaging mass spectrometry further demonstrated region-specific alterations in retinal phospholipid species with PEDF/PEDF-R deficiency, including DHA- and AA-containing phospholipids enriched within photoreceptor-associated layers, in particular, and in some inner retina regions. These findings reveal region-specific alterations in retinal phospholipid distribution associated with disruption of PEDF/PEDF-R signaling, providing direct in vivo evidence that PEDF/PEDF-R signaling regulates the regional distribution of retinal phospholipid species. Together, these findings support a model in which PEDF/PEDF-R signaling coordinates phospholipid remodeling with lipid droplet dynamics in photoreceptors and the RPE, to maintain the continuous renewal of photoreceptor outer segment membranes. The altered localization of TIP47 and accumulation of PLIN5-positive lipid droplets observed in mutant retinas are consistent with impaired fatty acid mobilization and disrupted membrane lipid turnover. Disruption of this axis leads to lipid dysregulation and, ultimately, photoreceptor cell death.

The progressive nature of photoreceptor degeneration in PEDF/PEDF-R-deficient retinas is supported by phosphatidylserine externalization, DNA fragmentation, and progressive disruption of outer segment architecture. Because photoreceptor outer segments undergo continuous membrane renewal, efficient phospholipid remodeling within photoreceptors together with phagocytic processing of shed outer segments by the RPE are essential for maintaining membrane integrity and retinal function. The accumulation of lipid droplets, altered phospholipid distribution, and structural abnormalities observed in mutant retinas suggest impaired lipid handling at the photoreceptor–RPE interface. Similar pathological features have been reported in other models of progressive retinal degeneration, including RPE65⁻/⁻ and *Rho^P23H/+^,* in which disruptions in outer segment processing, membrane integrity or lipid homeostasis contribute to photoreceptor loss. Taken together, our findings are consistent with a role for PEDF/PEDF-R signaling in supporting the phospholipid remodeling required for continuous outer segment renewal at the photoreceptor–RPE interface.

In addition to photoreceptor degeneration, PEDF/PEDF-R deficiency was associated with impaired retinal function and synaptic remodeling. Reduced ERG a-wave amplitudes indicate compromised photoreceptor responses, whereas diminished b-wave amplitudes suggest impaired synaptic transmission to downstream bipolar cells. Consistent with these functional deficits, rod bipolar cell dendrites exhibited reduced overlap with presynaptic terminals in the outer plexiform layer, indicative of dendritic remodeling at the first visual synapse. These findings indicate that PEDF/PEDF-R signaling is required to preserve both photoreceptor viability and synaptic organization within the outer retina.

The role of PEDF/PEDF-R signaling in lipid homeostasis may also have implications for retinal aging and disease. PEDF levels decline with age, and loss of PEDF or PEDF-R induces a senescence-like phenotype in the RPE (24–27). Because aging is accompanied by progressive changes in retinal lipid metabolism and is the major risk factor for age-related macular degeneration (AMD) (28), declining PEDF/PEDF-R signaling may contribute to lipid dysregulation, oxidative stress, and increased photoreceptor vulnerability.

In summary, our findings identify the PEDF/PEDF-R signaling axis as a previously unrecognized regulator of retinal phospholipid homeostasis, membrane integrity, and photoreceptor survival. By identifying a functional link between neurotrophic signaling and the phospholipid remodeling that supports photoreceptor membrane renewal, our findings provide a mechanistic framework for understanding how lipid metabolism supports photoreceptor survival and retinal function.

## Methods

### Animals

CRISPR-generated *Pnpla2* knockout mice on a C57BL/6J background were generated by the Genetic Engineering Core at the National Eye Institute (NEI) (8). *Serpinf1^+/−^* mice on a C57BL/6J background were described previously (11). Double-mutant mice were generated by crossbreeding *Pnpla2^+/-^* and *Serpinf1^+/-^* in the same C57BL/6J background.

All experiments were performed using 3-months-old mice of mixed sex, maintained under a 12-h light/12-h dark cycle with *ad libitum* access to standard chow and water. Studies were not restricted by sex, as no sex-specific differences in disease progression have been observed in these models. In accordance with the “Reduction” principle of the 3Rs, all bred animals were included in the study.

All procedures were approved by the NEI Animal Care and Use Committee and adhered to the Association for Research in Vision and Ophthalmology statement for the Use of Animals in Ophthalmic and Vision Research and the ARRIVE guidelines.

### RNA extraction, cDNA synthesis, and quantitative RT-PCR

Total RNA was isolated from mouse retinas using the RNeasy® Mini Kit (Qiagen, Germantown, MD) according to the manufacturer’s instructions. RNA (10-50 ng) was reverse-transcribed using the SuperScript III First-Strand Synthesis System (Thermo Fisher Scientific, Norristown, PA)

Quantitative real-time PCR was performed using a QuantStudio 7 Flex Real-Time PCR System (Thermo Fisher Scientific, Norristown, PA). *Pnpla*2 mRNA levels were normalized to *Hprt,* and *Serpinf1* mRNA levels to 18S rRNA. Reactions were performed using Taqman assays (Applied Biosystems, Pleasanton, CA) or the QuantiTect SYBR Green PCR kit (Qiagen, Germantown, MD). Primer sequences are listed in Table 1.

**Table 1:**
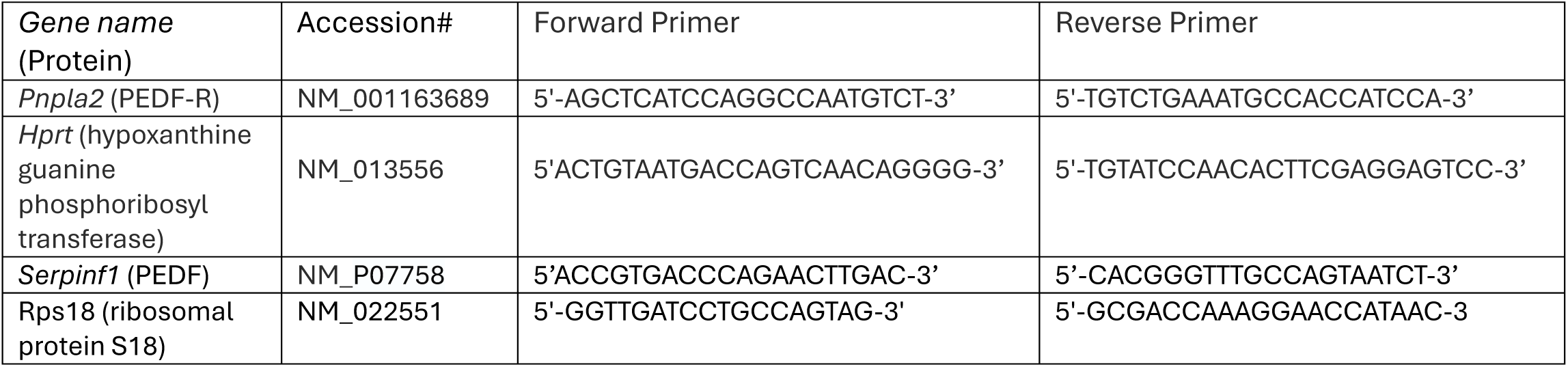
Primer Sequences.

### PEDF Enzyme-Linked Immunosorbent Assay (ELISA)

Protein extracts from retinas and RPE were prepared in RIPA Buffer (Thermo Fisher Scientific, Norristown, PA) supplemented with protease inhibitors (Pierce Protease Inhibitor Tablets, Thermo Fisher Scientific, Norristown, PA), using 80 μl of buffer per retina (8).

PEDF protein levels were quantified using a sandwich ELISA kit (XpressBio, Frederick, MD). Concentrations were calculated from a standard curve generated with purified PEDF (0– 10 ng/ml).

### Phospholipase A_1_ and A_2_ activity assay

PLA2 activities were measured using EnzChek™ Phospholipase A2 Assay Kit (Thermo Fisher Scientific, Norristown, PA). Retinal tissues were homogenized on ice using a sonic dismembrator (FisherBrand™ Model 50) (amplitude 30, 15 s, 4 °C), followed by centrifugation at 15,000 × *g* for 10 min at 4°C. The supernatants were used immediately.

Retinal extracts (50 µL) were incubated with substrate liposomes (50 µL) for 10 min at room temperature. Enzymatic activity was measured by ratiometric fluorescence (excitation 460 nm; emission 515/575 nm) using a SpectraMax® iD5 plate reader (Molecular Devices, Silicon Valley, CA). Bee venom PLA_2_ was used as standard for PLA_2_ activity, respectively.

### Histology and immunofluorescence

Eyes were enucleated, corneal incision made, and tissues fixed in 4% paraformaldehyde (Electron Microscopy Sciences, Morristown, PA) for ≥48 h. Samples were and paraffin-embedded. Ten-micrometer (10 µm) sections were cut from paraffin-embedded tissues and used for histology or Immunofluorescence.

Sections were incubated with primary antibodies (Table 2) followed by appropriate fluorescent secondary antibodies. Nuclei were counterstained with DAPI (0.1 μg/ml; Vector Laboratories, Newark, CA). Images were acquired using a ZEIS LSM 700 confocal microscope. Fluorescence intensity was quantified using ImageJ (14).

**Table 2:** Antibodies used in the study.

| Antibody | Type & host | Application | Dilution | Company,<br>Catalog Number |
| --- | --- | --- | --- | --- |
| Anti-Rhodopsin | Monoclonal<br>mouse | IF | 1:100 | Sigma, MAB5356 |
| Anti-Opsin blue | Polyclonal rabbit | IF | 1:200 | Sigma, AB-5407 |
| Anti-Synaptophysin | Monoclonal rabbit | IF | 1:250 | Abcam, Ab52636 |
| PKC alpha | Monoclonal<br>mouse | IF | 1:50 | Novus Biological,<br>NB600-201SS |
| Alexa Fluor 488 | Goat anti-Mouse<br>IgG (H+L) | IF | 1:200 | Thermo Fisher<br>Scientific, A-11001 |
| Alexa Fluor 555 | Goat anti-Rabbit<br>IgG (H+L) | IF | 1:200 | Thermo Fisher<br>Scientific, A-2142 |

Outer nuclear layer (ONL) thickness was measured on hematoxylin and eosin (HCE)-stained sections in defined retinal locations using ImageJ, and data were visualized as spider plots using GraphPad Prism.

### Transmission electron microscopy

Eyes were fixed in 2.5% glutaraldehyde in PBS (pH 7.4), post-fixed in 0.5% osmium tetroxide, and embedded in epoxy resin. Ultrathin sections (90 nm) were collected on 200-mesh copper grids, air-dried for 24 h, and double-stained with uranyl acetate and lead citrate. Images were acquired using a JEOL JM-1010 transmission electron microscope (8).

### Fundoscopy and detection of cell surface-exposed phosphatidylserine

Fluorescence fundoscopy was performed using a MICRON V retinal imaging system (Phoenix Research Labs, Pleasanton, CA) equipped with a 475/50 nm excitation filter (Semrock, Inc., Rochester, NY). PSVue-488 (Molecular Targeting Technologies, (West Chester, PA) was reconstituted at 1 mM in HBSS (Quality Biological) and administered topically as eye drops (5 μl per eye).

After 24 h, mice were anesthetized, pupils dilated with 1% tropicamide (Akorn, Lake Forest, Illinois) for 5 min, and eyes were lubricated with hypromellose gel (GenTeal) (0.3% hypomellose; Alcon, Fort Worth, TX) (14). Retinal images were acquired in vivo, and mean fundus fluorescence intensity was quantified using ImageJ.

### TUNEL assay

Photoreceptor cell death was assessed on paraffin-embedded retinal sections using the Click-iT Plus TUNEL assay kit (Invitrogen, Norristown, PA) according to the manufacturer’s instructions. Images were acquired using a Zeiss-LSM 700 confocal microscope.

### Imaging mass spectrometry and LC-MS/MS analysis

For IMS analysis, mouse eyes were sectioned at a thickness of 10 µm using a CM3050 Cryostat (LEICA CM3050S, Leica Microsystems Inc., Bannockburn, IL). Sections were thaw-mounted onto Indium-Tin-Oxide (ITO)-coated slides (Delta Technologies, Loveland, CO) and air-dried. 4-(dimethylamino) cinnamic acid (DMACA) was used for MALDI matrix in both positive and negative mode. Sublimation was performed at 175 °C for 15 min at a vacuum less than 200 mTorr using an in-house designed sublimation apparatus as described previously (29). MALDI MS data were acquired with a 10 μm pixel size using a timsTOF Pro MALDI imaging platform in qTOF mode with TIMS deactivated (Bruker Daltonik, Bremen, Germany). The mass spectrometer was calibrated with red phosphorus prior to data acquisition using a 120,000 FWHM (at m/z 400) mass resolving power. Data were acquired in either positive or negative ionization mode within a mass range of m/z 400 to 1500 with 100 laser shots per pixel. timsTOF data were loaded into SCiLS lab MVS (version 2024b Pro; Bruker Daltonics, Bremen, Germany). Features were extracted using Time-aligned-Region-complete-eXtraction feature finding with 1% relative intensity available in SCiLS Lab. Total ion current was used for normalization(30).

For lipid identification, two cryostat sections were collected and homogenized in PBS. Lipids were extracted using 2:2:1.8 methanol:chloroform:PBS. The samples were analyzed using a Vanquish ultrahigh performance liquid chromatography (UPLC) system interfaced to a Q Exactive HF mass spectrometer (Thermo Fisher Scientific, San Jose, CA) equipped with a HESI-II electrospray ionization source. Chromatographic separation was performed with a reverse-phase Acquity BEH C18 column (1.7 mm, 2.1×150mm, Waters, Milford, MA) at a flow rate of 250 ul/min and column temperature of 50°C under the following gradient: 0-1 min, 20% B(10:90 acetonitrile/isopropanol in 10 mM ammonium formate); 1-8 min: 20-100% B; 8–10 min, B = 100 % balanced with 40:60 H2O/acetonitrile in 10 mM ammonium formate. The mass spectrometer was operated in a data-dependent mode consisting of MS1 acquisition (R=60,000) from m/z 300-1600, using an MS AGC target value of 3E6, maximum ion time of 100 ms followed by up to 10 MS/MS scans (R=15,000) of the most abundant ions detected in the preceding MS scan. The MS2 AGC target value was set to 1E5 with a maximum ion time of 100 ms, and a normalized collision energy of 15, 30, 50, and dynamic exclusion was set to 7.5 s. Lipid identification was performed with MS-DIAL (version 5.5.250820) in lipidomics mode (30)

### Electroretinography (ERG)

ERG was performed using an Espion E2 system with a Color Dome stimulator (Diagnosys LLC, Lowell, MA). Mice were dark-adapted overnight and anesthetized under dim red light. Recording electrodes were placed on the cornea, a reference electrode in the mouth, and a ground electrode subdermally. Eyes were kept hydrated with GenTeal eye gel throughout the procedure. Scotopic responses were elicited using a series of flashes (15 flashes) ranging from 0.0001 to 10 cd.s/m^2^.

A-wave and b-wave amplitudes were measured from baseline measured and averaged across both eyes of each animal. Data were exported to Microsoft Excel for analysis. Both eyes were averaged to obtain amplitude values for each mouse.

### Experimental design and statistical analysis

Biological replicates were defined as individual animals, with one retina analyzed per animal unless otherwise specified. For histological analyses, ONL thickness was measured at defined distances from the optic nerve head in a single section per retina, and values were averaged across retinas within each group. For immunofluorescence experiments, 3–5 images per retina were acquired and quantified using ImageJ, and the mean value per retina was used as a single biological replicate. Data were analyzed using GraphPad Prism version 10.1.2. Comparisons between groups were performed using unpaired two-tailed Student’s *t* tests, and multiple-group comparisons were performed using one-way ANOVA with Dunette’s post hoc test, comparing each experimental group to the control.

All data are presented as mean ± SD. *P* < 0.05 was considered statistically significant.

## Acknowledgements

This research was supported in part by the Intramural Research Program of the National Institutes of Health (NIH), the National Eye Institute. The contributions of the NIH authors were made as part of their official duties as NIH federal employees and are in compliance with agency policy requirements and are considered Works of the United States Government. However, the findings and conclusions presented in this paper are those of the authors and do not necessarily reflect the views of the NIH or the U.S. Department of Health and Human Services.

## References

1. S. J. Fliesler, R. E. Anderson, Chemistry and metabolism of lipids in the vertebrate retina. Prog Lipid Res 22, 79–131 (1983).

2. S. P. Becerra, Mechanistic insights into the role of PEDF-R (PNPLA2) in photoreceptors. Biosci Rep 46 (2026).

3. L. Notari et al., Identification of a lipase-linked cell membrane receptor for pigment epithelium-derived factor. J Biol Chem 281, 38022–38037 (2006).

4. F. R. Steele, G. J. Chader, L. V. Johnson, J. Tombran-Tink, Pigment epithelium-derived factor: neurotrophic activity and identification as a member of the serine protease inhibitor gene family. Proc Natl Acad Sci U S A 90, 1526–1530 (1993).

5. T. L. Pham et al., Defining a mechanistic link between pigment epithelium-derived factor, docosahexaenoic acid, and corneal nerve regeneration. J Biol Chem 292, 18486–18499 (2017).

6. P. Subramanian et al., Pigment epithelium-derived factor (PEDF) prevents retinal cell death via PEDF Receptor (PEDF-R): identification of a functional ligand binding site. J Biol Chem 288, 23928–23942 (2013).

7. D. Lewandowski et al., Dynamic lipid turnover in photoreceptors and retinal pigment epithelium throughout life. Prog Retin Eye Res 89, 101037 (2022).

8. A. Bernardo-Colon et al., Ablation of pigment epithelium-derived factor receptor (PEDF-R/Pnpla2) causes photoreceptor degeneration. J Lipid Res 64, 100358 (2023).

9. J. Bullock et al., Degradation of Photoreceptor Outer Segments by the Retinal Pigment Epithelium Requires Pigment Epithelium-Derived Factor Receptor (PEDF-R). Invest Ophthalmol Vis Sci 62, 30 (2021).

10. M. Hara et al., PNPLA2 mobilizes retinyl esters from retinosomes and promotes the generation of 11-cis-retinal in the visual cycle. Cell Rep 42, 112091 (2023).

11. S. Dixit et al., PEDF deficiency increases the susceptibility of rd10 mice to retinal degeneration. Exp Eye Res 198, 108121 (2020).

12. A. F. Goldberg, O. L. Moritz, D. S. Williams, Molecular basis for photoreceptor outer segment architecture. Prog Retin Eye Res 55, 52–81 (2016).

13. T. D. Lamb, S. P. Collin, E. N. Pugh, Jr., Evolution of the vertebrate eye: opsins, photoreceptors, retina and eye cup. Nat Rev Neurosci 8, 960–976 (2007).

14. A. Bernardo-Colon et al., H105A peptide eye drops promote photoreceptor survival in murine and human models of retinal degeneration. Commun Med (Lond) 5, 81 (2025).

15. F. Mazzoni et al., Non-invasive in vivo fluorescence imaging of apoptotic retinal photoreceptors. Sci Rep 9, 1590 (2019).

16. A. V. Bulankina et al., TIP47 functions in the biogenesis of lipid droplets. J Cell Biol 185, 641–655 (2009).

17. A. R. Kimmel, C. Sztalryd, Perilipin 5, a lipid droplet protein adapted to mitochondrial energy utilization. Curr Opin Lipidol 25, 110–117 (2014).

18. B. Qiu, M. C. Simon, BODIPY 493/503 Staining of Neutral Lipid Droplets for Microscopy and Quantification by Flow Cytometry. Bio Protoc 6 (2016).

19. F. Polato, S. P. Becerra, Pigment Epithelium-Derived Factor, a Protective Factor for Photoreceptors in Vivo. Adv Exp Med Biol 854, 699–706 (2016).

20. L. W. Rowe et al., Gene-Agnostic Therapeutic Strategies for Inherited Retinal Diseases: Neuroprotection and Immunomodulation. Genes (Basel) 17 (2026).

21. Q. Huang, S. Wang, C. M. Sorenson, N. Sheibani, PEDF-deficient mice exhibit an enhanced rate of retinal vascular expansion and are more sensitive to hyperoxia-mediated vessel obliteration. Exp Eye Res 87, 226–241 (2008).

22. H. Shindou et al., Docosahexaenoic acid preserves visual function by maintaining correct disc morphology in retinal photoreceptor cells. J Biol Chem 292, 12054– 12064 (2017).

23. N. G. Bazan, Cellular and molecular events mediated by docosahexaenoic acid-derived neuroprotectin D1 signaling in photoreceptor cell survival and brain protection. Prostaglandins Leukot Essent Fatty Acids 81, 205–211 (2009).

24. L. Cao et al., Polarized retinal pigment epithelium generates electrical signals that diminish with age and regulate retinal pathology. J Cell Mol Med 22, 5552–5564 (2018).

25. A. M. Kolomeyer, I. K. Sugino, M. A. Zarbin, Characterization of conditioned media collected from cultured adult versus fetal retinal pigment epithelial cells. Invest Ophthalmol Vis Sci 52, 5973–5986 (2011).

26. A. M. Kolomeyer, I. K. Sugino, M. A. Zarbin, Characterization of conditioned media collected from aged versus young human eye cups. Invest Ophthalmol Vis Sci 52, 5963–5972 (2011).

27. J. J. Steinle, S. Sharma, V. C. Chin, Normal aging involves altered expression of growth factors in the rat choroid. J Gerontol A Biol Sci Med Sci 63, 135–140 (2008).

28. W. L. Wong et al., Global prevalence of age-related macular degeneration and disease burden projection for 2020 and 2040: a systematic review and meta-analysis. Lancet Glob Health 2, e106–116 (2014).

29. D. Anderson, Messinger, J. D., Allen, J., Djambazova, K. V., Kruse, A., Curcio, C. A., Schey, K. L., Caprioli, R. M., Spraggins, J., & Dufrense, M., atrix Sublimation via In-House Developed Sublimation Apparatus. . https://www.protocols.io/view/matrix-sublimation-via-in-house-developed-sublimat-cg8jtzun (2023).

30. H. Tsugawa, Ikeda, K., Takahashi, M., Satoh, A., Mori, Y., Uchino, H., Okahashi, N., Yamada, Y., Tada, I., Bonini, P., Higashi, Y., Okazaki, Y., Zhou, Z., Zhu, Z.-J., Koelmel, J., Cajka, T., Fiehn, O., Saito, K., Arita, M., & Arita, M, A lipidome atlas in MS-DIAL 4. Nature Biotechnology 38, 1159–1163. (2020).

