## Supplemental figures for "PEDF/PEDF-R Signaling Regulates Retinal Phospholipid Homeostasis and Photoreceptor Survival"

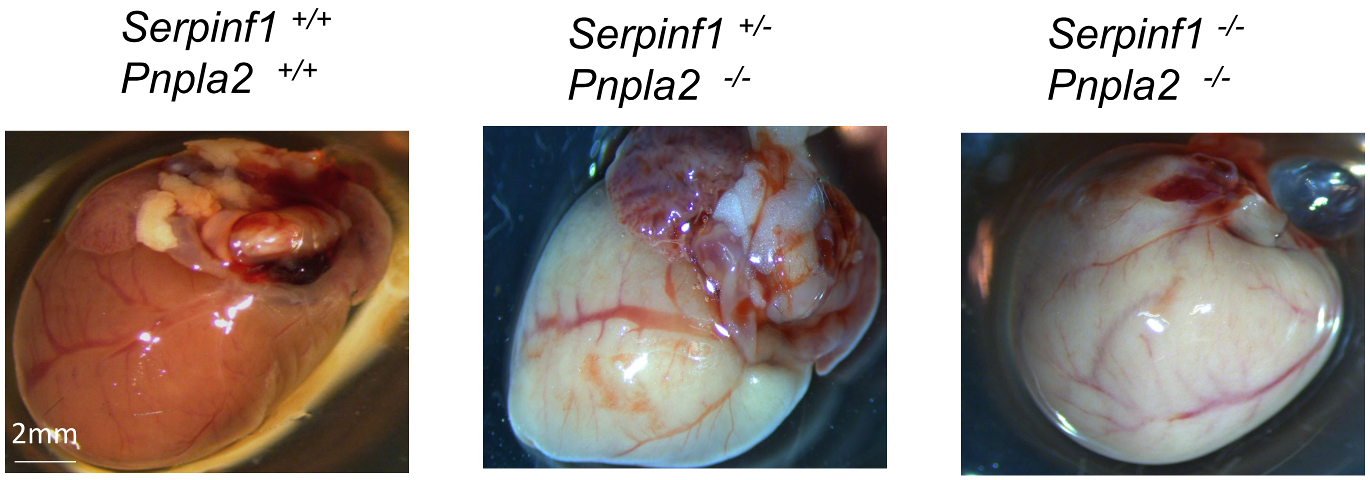


**Figure S1: Anatomical assessment of heart.**
Representatie images of hearts from *Serpinf1*$^{+/+}$ ; *Pnpla2*$^{+/+}$*, Serpinf1*$^{-/-}$ ; *Pnpla2*$^{-/-}$ and *Serpinf1*$^{+/-}$ ; *Pnpla2*$^{-/-}$ mice at 3 months of age. Hearts from *Serpinf1*$^{-/-}$ ; *Pnpla2*$^{-/-}$ and *Serpinf1*$^{+/-}$ ; *Pnpla2*$^{-/-}$ mice appeared en  and paler than those from *Serpinf1*$^{+/+}$ ; *Pnpla2*$^{+/+}$ controls.

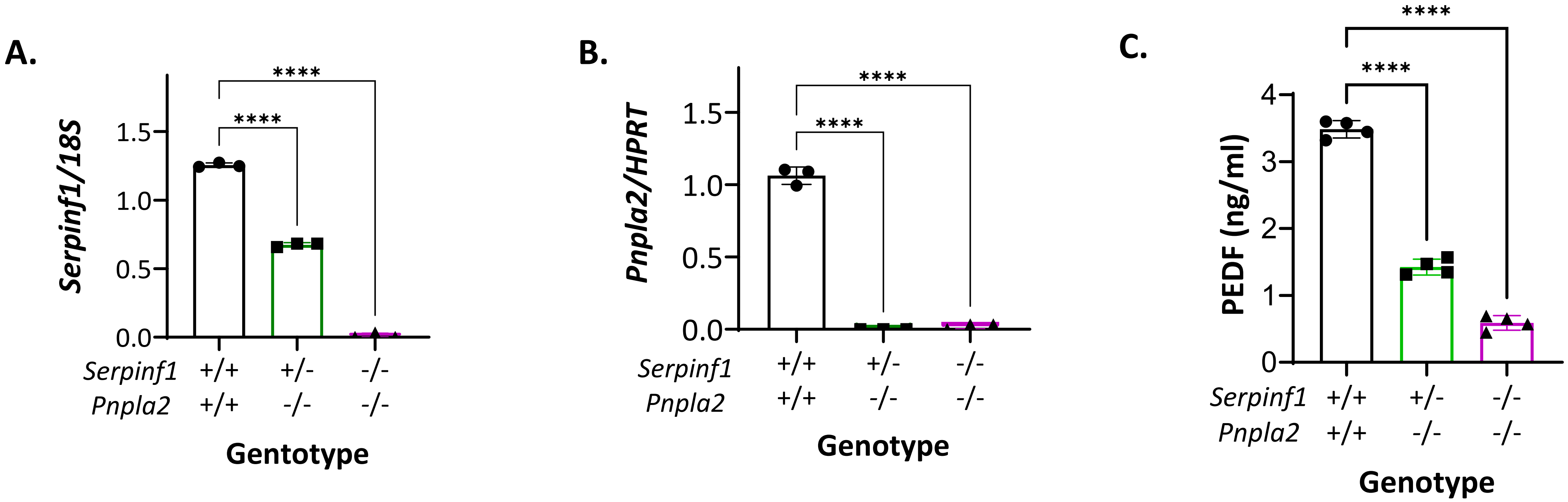


**Figure S2: Expression of *Serpinf1* (PEDF) and *Pnpla2* (PEDF-R) in the mouse retina.**

***(A, B)*** RT-PCR showing that *Serpinf1*$^{+/-}$ ; *Pnpla2*$^{-/-}$ and *Serpinf1*$^{-/-}$ ; *Pnpla2*$^{-/-}$ mice exhibited decreased and undetectable retinal *Serpinf1* gene expression, respectively (**A**), and both genotypes lacked *Pnpla2* gene expression relative to their *Serpinf1*$^{+/+}$ ; *Pnpla2*$^{+/+}$ (WT) control (**B**).

(**C**) ELISA showed that PEDF protein levels in RPE, the primary source of PEDF in the retina, was reduced in *Serpinf1*$^{+/-}$ ; *Pnpla2*$^{-/-}$ and *Serpinf1*$^{-/-}$ ; *Pnpla2*$^{-/-}$ mice compared with *Serpinf1*$^{+/+}$ ; *Pnpla2*$^{+/+}$ controls.

For RT-PCR, three retinas per genotype were analyzed, with each retina sample assay in triplicates. Each data point represents the average ± SD of the triplicate measurements for an individual retina. Statistical analysis was performed using one-way ANOVA followed by Dunnett's multiple-comparison test. For both mutant genotypes, ****$P < 0.0001$ relative to the 18S or HPRT housekeeping gene, as indicated in the y-axis.

For PEDF protein measurements, four RPE/choroid samples per genotype were analyzed in duplicate, with each data point representing the average ± SD of the duplicate measurements. A total of three mice (n=3) per genotype ere analyzed. Statistical analysis was performed using one-way ANOVA followed by Dunnett's multiple-comparison test. For both genotypes, ****$P < 0.0001$ relative to he *Serpinf1*$^{+/+}$ x *Pnpla2*$^{+/+}$ control.

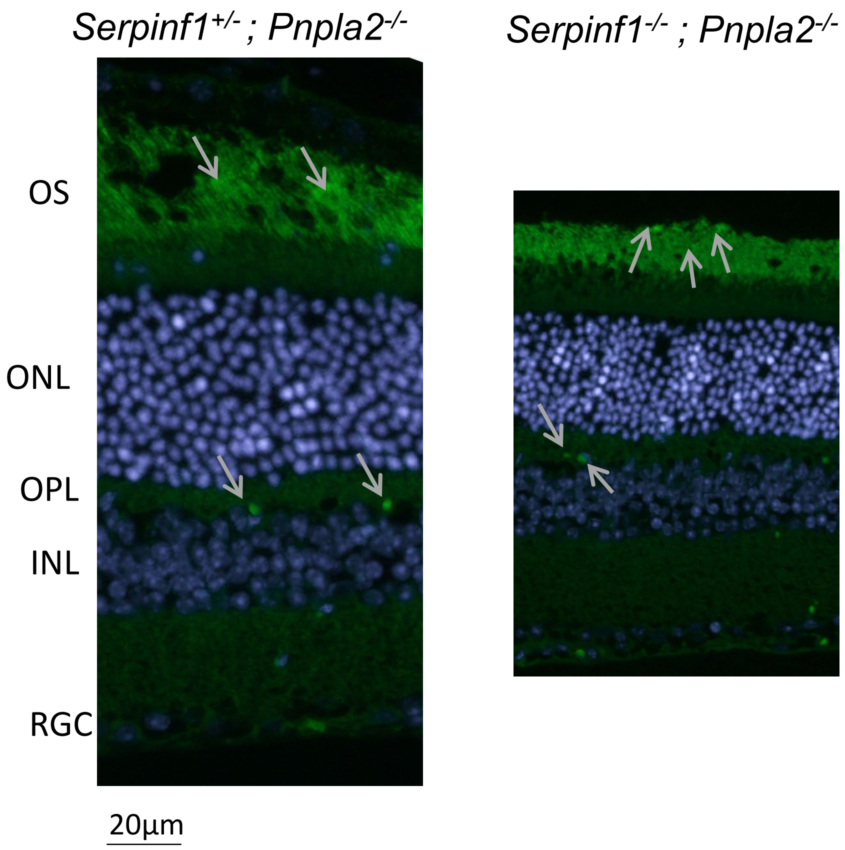


**Figure S3. PSVue labeling of phosphatidylserine (PS) externalization associated with photoreceptor cell death in *Serpinf1-* and *Pnpla2-d*eficient mice.**
Representative fluorescent micrographs of retinal cross-sections from *Serpinf1*$^{+/-}$ *; Pnpla2*$^{-/-}$ and *Serpinf1 -/- ; Pnpla2*$^{-/-}$ mice treated with 1 mM PSVue 488 eye drops at 3 months of age. PSVue 488 labeling (green) indicates phosphatidylserine (PS) externalization localized to the outer nuclear layer (ONL). Nuclei were counterstained with DAPI (blue). Four retinas per genotype (n=4) were analyzed. Scale bar = 20 µm.

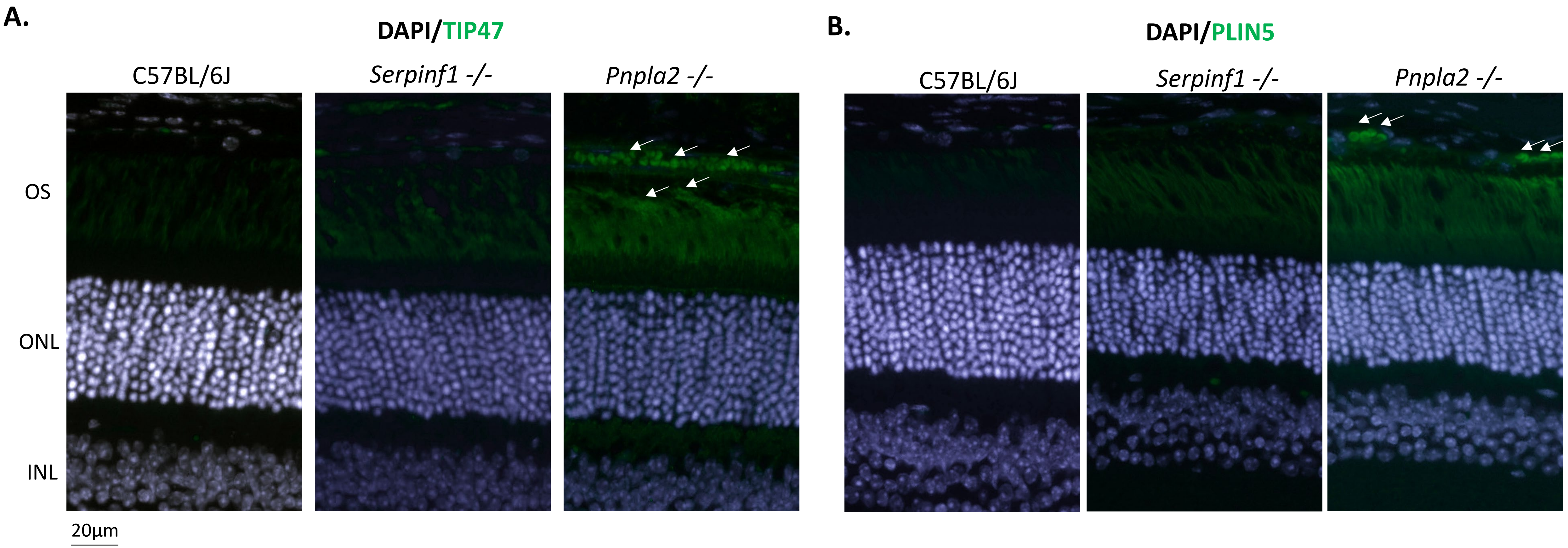


**Figure S4. Loss of PEDF does not result in lipid droplet accumulation in the retina.**
Representative fluorescence micrographs of retinas from 3-month-old C57BL/6J, *Serpinf1*$^{-/-}$, and *Pnpla2*$^{-/-}$ mice stained for TIP47 (tail-interacting protein of 47 kDa), a lipid droplet-associated protein, and PLIN5 (perilipin 5), a lipid droplet protein that promotes interactions between lipid droplets and mitochondria.

(A) TIP47 immunoreactivity was detected in the retinal pigment epithelium (RPE)/choroid and photoreceptor outer segments (OS; white arrows) of *Pnpla2*$^{-/-}$ mice, indicating lipid droplet accumulation. In contrast, no obvious lipid droplet accumulation was observed in *Serpinf1*$^{-/-}$ retinas.

(B) PLIN5 staining was prominent in the RPE of *Pnpla2*$^{-/-}$ mice (white arrows), consistent with lipid droplet accumulation, whereas little to no PLIN5 staining was detected in *Serpinf1*$^{-/-}$ retinas. For all immunofluorescence analyses, retinas from three animals per genotype were examined.

(C) Scale bar = 20 µm.

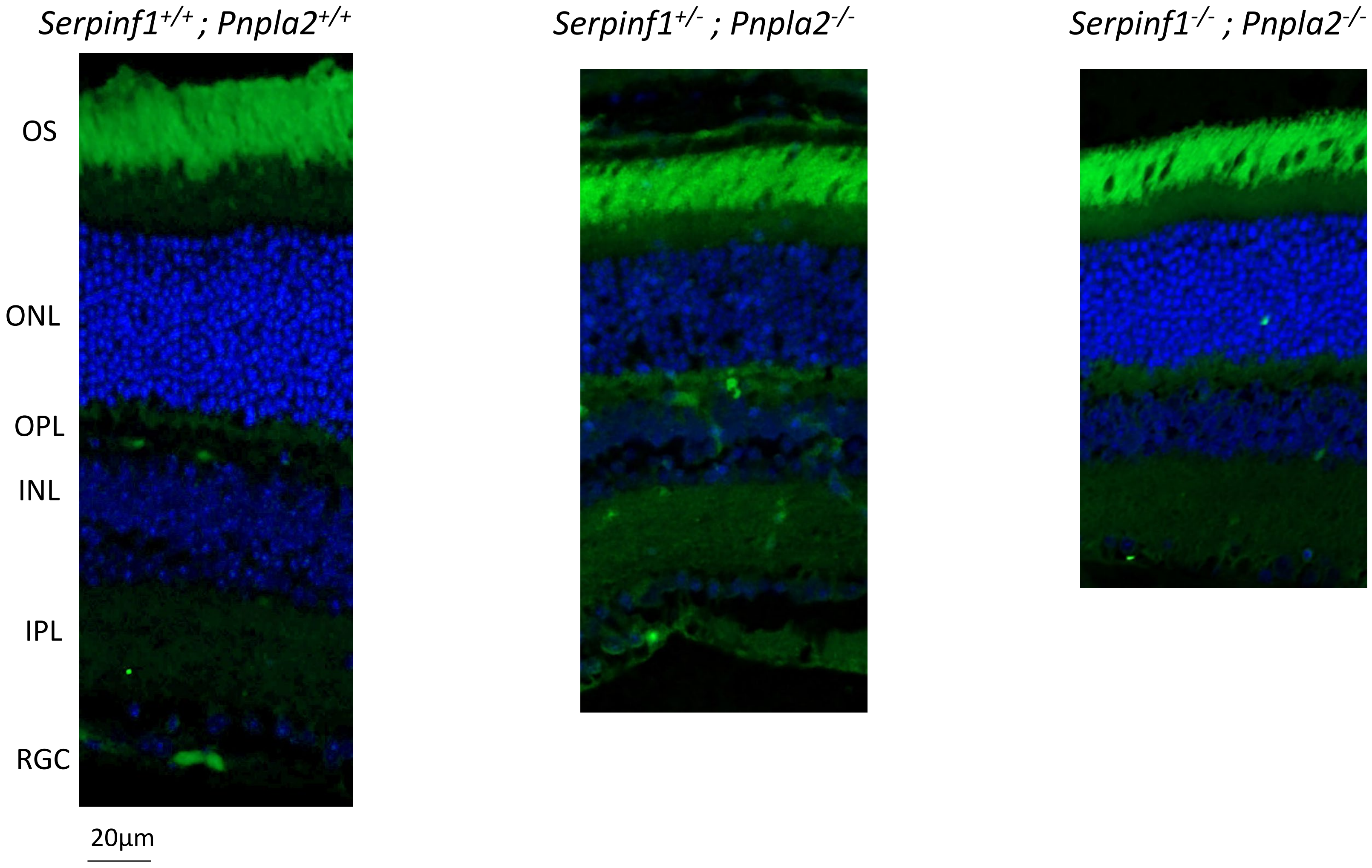


**Figure S5. Isolectin B4 staining reveals normal retinal vascular patterning in *Serpinf1*- and *Pnpla2*-deficient mice:** Representative fluorescence micrographs of retinal sections from *Serpinf1*$^{-/-}$; *Pnpla2*$^{-/-}$, *Serpinf1*$^{+/-}$; *Pnpla2*$^{-/-}$,and *Serpinf1*$^{+/+}$; *Pnpla2*$^{+/+}$ mice at 3 months of age. Retinal vasculature was visualized by isolectin B4 (IB4) staining, and nuclei were counterstained with DAPI. No evidence of retinal neovascularization or overt alterations in vascular patterning was observed among the genotypes examined. Three retinas per genotype were analyzed. Scale bar, 20 µm.
